# Individual variation underlying brain age estimates in typical development

**DOI:** 10.1101/2020.11.30.405290

**Authors:** Gareth Ball, Claire E Kelly, Richard Beare, Marc L Seal

## Abstract

Typical brain development follows a protracted trajectory throughout childhood and adolescence. Deviations from typical growth trajectories have been implicated in neurodevelopmental and psychiatric disorders. Recently, the use of machine learning algorithms to model age as a function of structural or functional brain properties has been used to examine advanced or delayed brain maturation in healthy and clinical populations. Termed ‘brain age’, this approach often relies on complex, nonlinear models that can be difficult to interpret. In this study, we use model explanation methods to examine the cortical features that contribute to brain age modelling on an individual basis.

In a large cohort of n=768 typically-developing children (aged 3-21 years), we build models of brain development using three different machine learning approaches. We employ SHAP, a model-agnostic technique to estimate sample-specific feature importance, to identify regional cortical metrics that explain errors in brain age prediction. We find that, on average, brain age prediction and the cortical features that explain model predictions are consistent across model types and reflect previously reported patterns of regional brain development. However, while several regions are found to contribute to brain age prediction, we find little spatial correspondence between individual estimates of feature importance, even when matched for age, sex and brain age prediction error. We also find no association between brain age error and cognitive performance in this typically-developing sample.

Overall, this study shows that, while brain age estimates based on cortical development are relatively robust and consistent across model types and preprocessing strategies, significant between-subject variation exists in the features that explain erroneous brain age predictions on an individual level.

## Introduction

Human brain development involves dynamic and interdependent changes that occur in a protracted manner throughout childhood and adolescence [Amlien et al., 2016; Giedd and Rapoport, 2010; Lyall et al., 2015]. Magnetic resonance imaging (MRI) has provided an unprecedented method for studying typical brain developmental trajectories *in vivo*. Longitudinal MRI studies have revealed the differential timing of developmental trajectories across different brain tissue types and regions. Cortical thickness and surface area increase rapidly after birth [Lyall et al., 2015; Wang et al., 2019]. Subsequently, the majority of studies have demonstrated decreases in cortical thickness with age over childhood and adolescence, along with childhood increases followed by subtle adolescent decreases in cortical surface area [Brown et al., 2012; Ducharme et al., 2016; Tamnes et al., 2017]. Detailed understanding of these typical cortical developmental trajectories is critical, given deviations from typical trajectories have been linked to variations in cognitive performance [Burgaleta et al., 2014; Shaw et al., 2006], and implicated in neurodevelopmental and psychiatric disorders such as autism spectrum disorder, attention deficit hyperactivity disorder, and schizophrenia [Baribeau and Anagnostou, 2013; Mensen et al., 2016; Shaw et al., 2007].

MRI yields high-dimensional datasets characterised by many correlated features (e.g.: area, volume or cortical thickness of brain regions, voxels, or vertices), often with complex interactions. Generally, the number of features is greater than the number of subjects in the study (p >> n), limiting the use of traditional statistical techniques. Modern machine learning approaches are well suited to tackling such problems, with examples focused on grouping together related regions and/or metrics to reduce the number of features through dimension reduction [Ball et al., 2019; Beckmann and Smith, 2005; Groves et al., 2012; Sotiras et al., 2015; Sotiras et al., 2017; Xu et al., 2009], using kernels to combine information across multiple features [Dyrba et al., 2015; Klöppel et al., 2008; Mourão-Miranda et al., 2005; Rondina et al., 2018] and employing sparse or highly-regularised methods that penalise complex models, thus limiting the potential to overfit to noise in large-p datasets [Baldassarre et al., 2017; Beer et al., 2019; Lin et al., 2014; Rasmussen et al., 2012; Teipel et al., 2017]. In large study populations, deep learning approaches have also proven effective [Dinsdale et al., 2021; Jonsson et al., 2019] although recent evidence suggests that these models offer relatively minimal performance gain in brain MRI studies compared to simpler methods [He et al., 2020; Schulz et al., 2020b].

The applications of machine learning to MRI data generally fall into two categories: classification, e.g.: the grouping of subjects based on disease status, or the segmentation of images based on tissue characteristics; and regression, e.g: the prediction of subject characteristics (for example, disease severity or cognitive performance) based on properties derived from brain MRI. A recent application that has gained popularity is the prediction of subject age from brain MRI, termed ‘brain age’ [Cole and Franke, 2017; Franke and Gaser, 2019]. In this setting, age is modelled as a function of structural or functional brain properties in a large training sample with parameters from this model then applied to unseen data to predict age. The discrepancy between the age predicted with reference to the normative model and the true chronological age (i.e., the brain age delta or brain age gap) can be used to index the extent to which individuals deviate from a typical developmental or ageing trajectory [Cole et al., 2015; Franke and Gaser, 2019].

Brain age modelling has most commonly been applied in adult populations, where a positive brain age gap (indicating the brain was predicted to be older than the chronological age) has been associated with advanced aging in many disorders, including schizophrenia, major depressive disorder, traumatic brain injury, and Alzheimer’s disease and mild cognitive impairment [Cole et al., 2015; Franke and Gaser, 2012; Han et al., 2020; Kaufmann et al., 2019; Koutsouleris et al., 2014]. Studies in children are less frequent and their findings are more variable. In typically developing children, studies have observed both positive [Erus et al., 2015; Zhao et al., 2019] and negative associations between brain age gap and cognitive performance in development [Lewis et al., 2018]. Others have reported very small, or nonsignificant, effect sizes for the association between brain age gap and cognitive performance [Ball et al., 2017; Khundrakpam et al., 2015]. In children with autism spectrum disorder, a negative brain age gap (reflecting that the brain was predicted to be younger than expected for chronological age) was related to greater disorder symptom severity [Tunç et al., 2019]. In a non-typically developing sample of children and adolescents, a positive brain age gap was associated with greater psychopathology symptom severity [Cropley et al., 2020].

Variable findings in paediatric studies may occur because, while deviation is typically thought of as an acceleration or deceleration of ongoing developmental processes, brain age modelling projects many complex, non-linear and potentially overlapping patterns of brain development into a single summary metric. Studies have also demonstrated that factors unrelated to brain development or aging can affect brain age estimates [Franke et al., 2015; Le et al., 2018b] with differences in imaging metrics, scanner acquisitions, models and model parameters all contributing to brain age prediction accuracy [Brouwer et al., 2020; Franke and Gaser, 2012; Han et al., 2020; Valizadeh et al., 2017; Varikuti et al., 2018]. It is vital to identify the factors that underlie individual brain age prediction estimates in order to understand how deviations in neuroanatomical trajectories may relate to neurodevelopment in both healthy and clinical populations.

Models employed in brain age modelling are often nonlinear, so called ‘black boxes’, making it difficult to interpret or determine which features contribute to model predictions [Cole and Franke, 2017; Jiang et al., 2020; Liem et al., 2017]. Other methods, such as regularised linear regressions, return model coefficients that may be used directly to understand features that drive predictions on a group average level [Khundrakpam et al., 2015; Niu et al., 2020]. However, the contribution of different features to brain age estimates may vary over time [Brown et al., 2012], and across individuals. Several methods have been proposed to allow model-agnostic explanation or interpretation of machine learning models [Bach et al., 2015; Chen et al., 2018; Lundberg et al., 2020; Montavon et al., 2018; Ribeiro et al., 2016; Shrikumar et al., 2019; Štrumbelj and Kononenko, 2014]. A recent addition, Shapley Additive Explanations (SHAP), presents a unified framework for estimating model-agnostic feature importance on a sample-by-sample basis that is equally applicable across linear, nonlinear and deep models [Lundberg et al., 2020; Lundberg and Lee, 2017]. SHAP fits local models to small sections of the decision space, synthesising new datapoints by sampling values for a single feature from a background dataset and assigning importance based on game-theoretic principles [Lundberg and Lee, 2017]. In MRI studies, SHAP can be used to estimate the brain features (regions, voxels, or vertices) that contribute to a model’s prediction for a given subject. This permits further examination of the similarities and differences across subjects, clustering or classifying individuals based on how they are perceived by the model [Schulz et al., 2020a].

In this study, we combine brain age modelling with model explanation to explore how individual variation in neuroanatomy drives brain age prediction in a large cohort of typically-developing children and adolescents (n=768). Across three different model types (regularised linear regression, nonlinear Gaussian process regression and tree-ensemble regression), we show that brain age estimates are generally correlated across models, as are the important features that underlie each model’s predictions *on average*. Using fine-grained subject-by-subject model explanations, we find feature-level contributions to brain age prediction are distributed across the cortex and vary significantly across individuals, even when subject age, sex and brain age predictions are similar. In this study sample, we also find no evidence of an association between cognitive performance and the morphological changes underlying brain age estimation in typical development.

## Materials and Methods

### Participants

Data were acquired from the Pediatric Imaging, Neurocognition and Genetics (PING) Study [Jernigan et al., 2016] with participants from several US sites across a wide age and socioeconomic range. The human research protections programs and institutional review boards at all institutions participating in the PING study approved all experimental and consenting procedures, and all methods were performed in accordance with the relevant guidelines and regulations [Brown et al., 2012]. Written parental informed consent was obtained for all PING subjects below the age of 18 and directly from all participants aged 18 years or older. Exclusion criteria included: a) neurological disorders; b) history of head trauma; c) preterm birth (less than 36 weeks); d) diagnosis of an autism spectrum disorder, bipolar disorder, schizophrenia, or significant intellectual disability; e) pregnancy; and f) daily illicit drug use by the mother for more than one trimester [Jernigan et al., 2016]. Data from the PING study are made available via the NIMH Data Archive (https://nda.nih.gov/edit_collection.html?id=2607).

The PING cohort included 1493 participants aged 3–21 years; of these, neuroimaging data for n = 773 were available to download (March 2016). After quality control and image processing (see below), the final cohort comprised n = 768 participants (mean [S.D] age = 12.28 [5.02] years; 404 male) acquired from seven study scanners/sites. Site/scanner-specific demographic data are shown in Table S1.

### Neuroimaging data

3 Tesla, T1-weighted images were acquired using standardized high-resolution 3D RF-spoiled gradient echo sequence with prospective motion correction (PROMO), with pulse sequences optimized for equivalence in contrast properties across scanner manufacturers (GE, Siemens, and Phillips) and models (for details, see [Jernigan et al., 2016]) Site-specific imaging parameters are shown in Table S2.

### Image processing

Structural T1 volumes were processed as described previously [Ball et al., 2017; Ball et al., 2020]. Briefly, vertex-wise maps of cortical thickness and area were constructed with FreeSurfer 5.3 (http://surfer.nmr.mgh.harvard.edu). This process includes removal of non-brain tissue, transformation to Talairach space, intensity normalisation, tissue segmentation and tessellation of the grey matter/white matter boundary followed by automated topology correction. Cortical geometry was matched across individual surfaces using spherical registration and maps smoothed to 10 mm FWHM [Dale et al., 1999; Fischl et al., 2002; Fischl and Dale, 2000]. To reduce computational burden, cortical data were downsampled to the *fsaverage5* surface comprising 10242 vertices per hemisphere.

Quality control assessment for the PING data is detailed in [Jernigan et al., 2016]. Images were inspected for excessive distortion, operator compliance, or scanner malfunction. Specifically, T1-weighted images were examined slice-by-slice for evidence of motion artefacts or ghosting and rated as acceptable, or recommended for re-scanning. We performed additional, on-site, visual quality control [Ball et al., 2017]. Any images that failed initial surface reconstruction, or returned surfaces with topological errors, were manually checked and resubmitted to FreeSurfer.

For all subjects, cortical thickness and area maps were parcellated into n = 179 labels per hemisphere using a parcellation scheme based on anatomical and functional cortical boundaries (excluding the hippocampus) [Glasser et al., 2016]. We repeated our analysis using an alternative parcellation comprising n = 250 regions per hemisphere with approximately equal area [Arnatkeviciūtė et al., 2019; Ball et al., 2020]. Regional measures of mean cortical thickness and surface area were calculated using the 95% trimmed mean and log-transformed sum of all vertex values contained within the assigned region, respectively.

### Brain age estimation

We employed three machine learning algorithms to estimate brain age: regularised linear regression, Gaussian process regression and an ensemble of gradient boosted regression trees, XGBoost, detailed below:

#### Linear model: Regularised linear regression

We fit a regularised linear regression model with elastic net penalty [Zou and Hastie, 2005]. This approach combines LASSO and Ridge regression, regularising both the ℓ_1_ and ℓ_2_-norms of the model coefficients to produce a sparse linear model with most coefficients set to zero and the variance of the remaining non-zero coefficients regularised. The elastic net minimises:

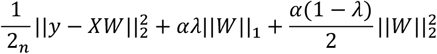

Where *W* = [*w*_1_ …, *w_p_*] represents the model coefficients, and ||*W*||_1_ and 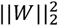 represent the ℓ_1_ and ℓ_2_ penalties, respectively. The parameters, *α* and *λ* are hyperparameters that regulate the strength and balance of the two penalty terms. Elastic net models are well suited to high-dimensional problems and have proven successful in brain age prediction [Khundrakpam et al., 2015; Niu et al., 2020]. Hyperparameters were tuned using nested cross-validation (see ‘Model training and evaluation’ below)

#### Nonlinear model: Gaussian process regression

Gaussian process regression (GPR) is a highly flexible, Bayesian modelling approach for the prediction of continuous variables [Rasmussen and Williams, 2006]. A major benefit of GPR is the ability to fit nonlinear functions without needing to specify the functional form (quadratic, cubic, etc.) or define any inflection points. GPR has previously been used successfully to model relationships between MRI-derived metrics and age [Ball et al., 2020; Cole et al., 2015; Cole et al., 2018]. We review GPR below; for a technical overview we refer interested readers to [Rasmussen and Williams, 2006].

A Gaussian Process is a distribution of functions that map: *x* → *y* and is fully specified by a mean function, *m* = *μ*(*x*) = 0 and a covariance function, *cov* = *k*(*x, x^T^*) such that *f* ~ *GP*(*m, cov*). The covariance function, *k*, models the dependence of values of *f* at different values of *x, x_i_* and *x_j_*, respectively. Prior to model fitting, we defined *k*(*x_t_, x_j_*) with a combined linear and radial basis function (RBF) kernel as:

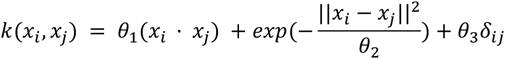

Where *δ* models observation noise when *i* == *j*, otherwise *δ* = 0, and *θ*_1,2,3_ are hyperparameters learned from the data by optimising the log-marginal-likelihood of the model during training.

#### Ensemble model: XGBoost

XGBoost is an ensemble model based on gradient tree boosting [Chen and Guestrin, 2016] that has been shown to accurately predict age from MRI [Kaufmann et al., 2019]. Gradient tree boosting builds a predictive model from an ensemble of weak base learners, often shallow regression trees. In contrast to other ensemble models, such as Random Forest, the model is trained in an additive manner with learners added to the model sequentially to correct prediction errors from previous model estimates. Gradient boosting methods are highly flexible and may be prone to overfit in smaller samples sizes. Regularisation is provided through a series of hyperparameters including the learning rate, tree depth, weight shrinkage and data subsampling. For each model, we chose a relatively low learning rate of 0.05 and optimised additional hyperparameters as set out in Table 1.

**Table 1:** Hyperparameter optimisation for XGBoost

| Hyperparameter | Distribution |
| --- | --- |
| Number of estimators | 100, 250, 500 |
| Features sampled by tree | 0.5 - 1.0 |
| Minimum child weight | 1, 2, 3, 4, 5 |
| Maximum tree depth | 2, 3, 4 |
| Subjects sampled each iteration | 0.5 - 1.0 |
| $\ell_2$ - regularisation | 10 - 100 |

### Model training and evaluation

All models were evaluated with 5-fold cross-validation, stratified by site (CV). Regional cortical thickness and area data were first concatenated to form an *n* × *p* matrix. Prior to model training, we performed ComBat harmonisation to account for potential site and scanner effects within the PING cohort [Fortin et al., 2018] and scaled data to unit variance and zero mean. In each fold, harmonisation and standardisation parameters were estimated in the training data, before being used to transform test samples. For linear (Elastic Net) and ensemble (XGBoost) models, hyperparameters were tuned using a randomised parameter search (n=500 iterations) within a nested 5-fold CV loop during training.

For each sample in the test set, brain age delta (also known as brain age gap) was calculated by subtracting age from predicted brain age. Model accuracy was evaluated using the median absolute error (MAE) of age prediction and the R^2^ score between true and predicted ages, averaged over the five folds.

All model training and evaluation was performed using *scikit-learn* (0.22.1) implemented in Python (3.8.3) [Pedregosa et al., 2011]. To fit the ensemble model, we used the *XGBoost* Python library (https://github.com/dmlc/xgboost) with scikit-learn API. ComBat harmonisation was performed using *neurocombat-sklearn* (https://github.com/Warvito/neurocombat_sklearn). Cortical surfaces are visualised using *pysurfer* (0.10) (https://github.com/nipy/PySurfer/). Model training and evaluation code is available from: https://github.com/garedaba/brainAges

### Alternative preprocessing

We repeated our analyses employing alternative preprocessing steps before model fitting: regressing out variance explained by mean cortical thickness and total surface area from regional measures; and employing a dimension reduction step with Principal Component Analysis (PCA; retaining 90% of the variance in the data).

### Model explanations

We employed Shapley Additive Explanations (SHAP) to estimate individual-level explanations for each model [Lundberg et al., 2020]. SHAP is a game-theoretic approach to additive feature attribution, where a change in model outcome with respect to a baseline (i.e.: the average prediction from the whole sample) can be apportioned to individual features, such that:

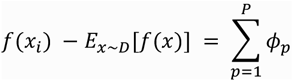

Where *f*(*x_i_*) represents the model output for a single sample; *D* represents the full training, or background, data set and *ϕ_p_* represents the feature importance, or SHAP value, of a single feature, p. The sum of importances across all features equals the difference between the predicted output and the expected model output given the background data. This is achieved by repeatedly fitting local linear regression models to subsets of features, synthesising new datapoints by sampling values for a single feature from the background data and assigning importance based on the degree to which the model output is changed. Exact importance values can be calculated directly for linear models with independent features as: 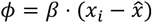 where *β* refers to the linear regression coefficients and 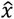 is the mean of the training, or background, data. Similarly, exact methods exist for tree-based models, for a technical overview, we refer readers to Lundberg et al. [Lundberg et al., 2020]. For consistency, we have used the kernel SHAP approach, which is model-agnostic and applicable to all three models used in this study.

As noted above, SHAP values are defined relative to a background reference set. This can be sampled from the whole training dataset, or chosen to reflect a different baseline. In Figure 1, we illustrate how brain age model explanations differ based on the choice of baseline. Using samples from the full dataset, feature importance sums to the difference between model output and the expected model output, *E*(*f*(*x*)), which in this case is the average participant age or approximately 12 years (Fig 1A). As such, positive values would denote where increases in thickness or area in important regions drives the model to predict an older (than 12) brain age, and vice versa. An alternative approach is to select age-matched background samples based on the age of each test subject. Here (Fig 1B) the nearest *k*-neighbours based on age in the training set are used to define the baseline, meaning that positive values of important features now denote how increases (alternatively decreases) in regional thickness or area drive the model to predict an older or younger brain age *relative to subjects with similar ages*.

**Figure 1:**
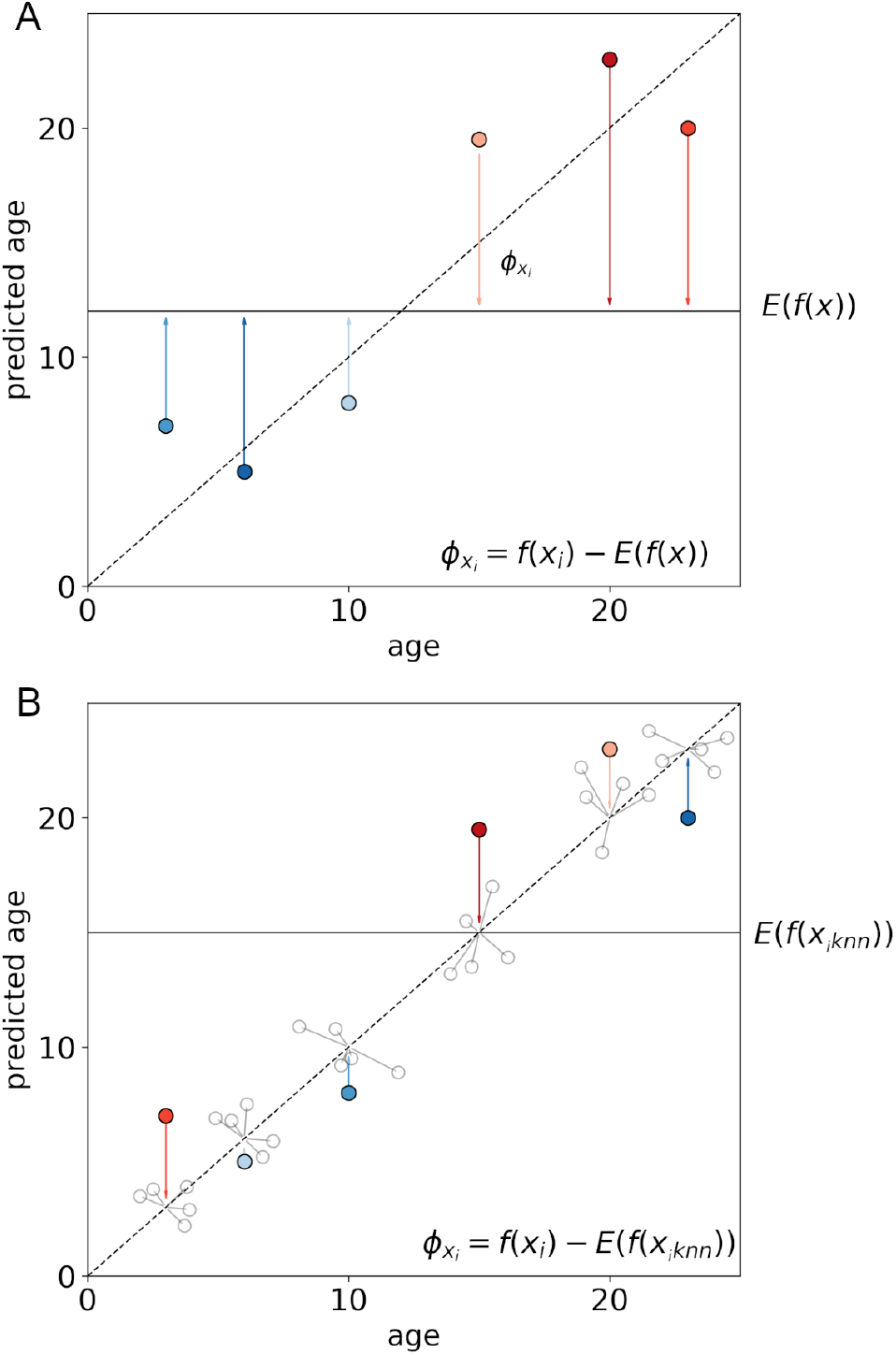
Calculating model explanations using different baselines. A. For a given dataset, the expected model output represents the sample mean, *E*(*f*(*x*)). For a given test sample, *x_i_*, the sum of feature importances, *ϕ_x_i__*, equals the difference between model output and the average model output. B. An alternative baseline, uses the expected outcome from the *k*-nearest neighbours of *x_i_* as the baseline, providing model explanations that better reflect deviations from true sample age.

Within each CV fold, we defined the background reference set using 10-nearest neighbours in the training sample to calculate age-specific feature importances for every test subject, for each model. This resulted in an *n*-subject by *p*-feature ‘model explanation’ matrix, where each column reflects the importance of a given region’s thickness or area to an individual’s brain age prediction, relative to the age-matched background samples (i.e.: brain age delta; Fig 1B). The kernel SHAP method applies ℓ_1_ regularisation to select the number of features in the final linear model used to explain each prediction. We set this to allow a maximum of 50% of cortical features. This resulted in between *n*=114 and *n*=358 features selected for each subject (linear model w/ HCP parcellation: mean [S.D.] *n* = 342.2 [17.2]; nonlinear = 348.8 [12.8]; ensemble = 344.9 [17.0]). We also repeated our analysis using a smaller, fixed number of features (*n*=50).

To calculate model explanations, we used the *SHAP* (0.35) package (https://github.com/slundberg/shap).

### Variance partitioning

To investigate the role of confounding variables on estimates of brain age error, we use linear regression models to partition variance explained in brain age delta by age, sex, mean cortical thickness and total cortical surface area [Dinga et al., 2020]. Using cross-validated predictions of brain age and corresponding model explanations, we decomposed variance in brain age delta into variance explained by confounding variables (age, sex, global cortical metrics) and predictors (model explanations) as:

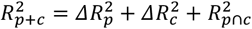

where 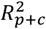 is the total variance explained in brain age delta by a linear model including all predictor and confounder terms; 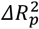 is the additional variance attributable to only predictor terms, 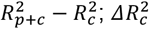 is the additional variance attributable to only confounder terms, 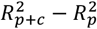 and 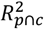 is the variance explained in brain age delta by factors shared by both predictors and confounders, 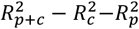 [Dinga et al., 2020]. To account for potential problems when fitting linear models to the high-dimensional model explanation matrix, we employed a Bayesian ridge regression model with a Gaussian prior over model coefficients. The regularization parameters were estimated during model fitting by maximizing the log marginal likelihood. Further, prior to model fitting, the model explanation matrix was projected to n=100 principal components, accounting for between 75% and 93% of variance in model explanations across the three models.

### Random surrogate maps

We compared inter-subject correspondence of feature importance maps to a null distribution constructed from a set of random surrogate maps with spatial autocorrelation and intensity distributions matched to the individual model explanations [Burt et al., 2020; Viladomat et al., 2014]. Surrogate maps were generated using BrainSMASH (https://brainsmash.readthedocs.io/en/latest/), creating 500 random maps for each subject, separately for each hemisphere and each feature set (regional thickness importance, regional area importance).

### Cognitive data

In addition to imaging data, participants in the PING study undertook comprehensive behavioural and cognitive assessments (NIH Toolbox Cognition Battery, NTCB). The NTCB comprises a set of tests that measure abilities across several cognitive domains, including cognitive flexibility, inhibitory control, and working memory, resulting in eight measures of cognitive performance (for full details of these measures in the PING cohort refer to: [Akshoomoff et al., 2014]). In total, neurocognitive data were available for n = 573 (75%) of the study sample.

To test associations between brain age delta and cognitive performance, we first modelled each raw cognitive score as a smooth function of age with an additional main effect of sex using a Generalised Additive Model (GAM), implemented by the *mgcv* package in *R* [Akshoomoff et al., 2014; Wood, 2017]. For each score, the residuals of this model represent individual performance at a level greater or lesser than expected for a given age and sex. Similarly, brain age delta, with variance due to confounding variables removed, represent brain age estimates greater or lesser than expected for a given age.

As performance across scales is closely correlated within individuals, and in order to avoid a large number of related statistical tests, we combined the standardised residuals of each of the cognitive models using PCA. This resulted in a set of components, each a linear combination of the original, residualised NTCB variables. We tested the association between brain age delta and the first two principal components of cognitive performance using linear regression.

## Results

### Accurate prediction of age from cortical metrics

We trained three machine learning models to predict age using regional measures of cortical thickness and area in a large cohort of typically-developing children and adolescents (n=768). Figure 2 illustrates model performance across all three brain age models: Elastic Net regression (‘linear’), Gaussian Process Regression (‘nonlinear’) and XGBoost (‘ensemble’). All three models achieved a high accuracy (mean MAE over crossvalidation folds [S.D.] = 1.81 [0.06], 1.75 [0.07], 1.92 [0.10]; R^2^ = 0.79 [0.01], 0.81 [0.01], 0.78 [0.03] for linear, nonlinear and ensemble models, respectively; Figure 2A, Figure 2B). Individual differences between predicted and true age, the estimated brain age delta, were also consistent across models (Figure 2C), with very high agreement between the regularised linear model and the nonlinear GPR model (r=0.96) and, to a lesser extent between linear and nonlinear models and the ensemble XGBoost model (r=0.81, 0.80).

**Figure 2:**
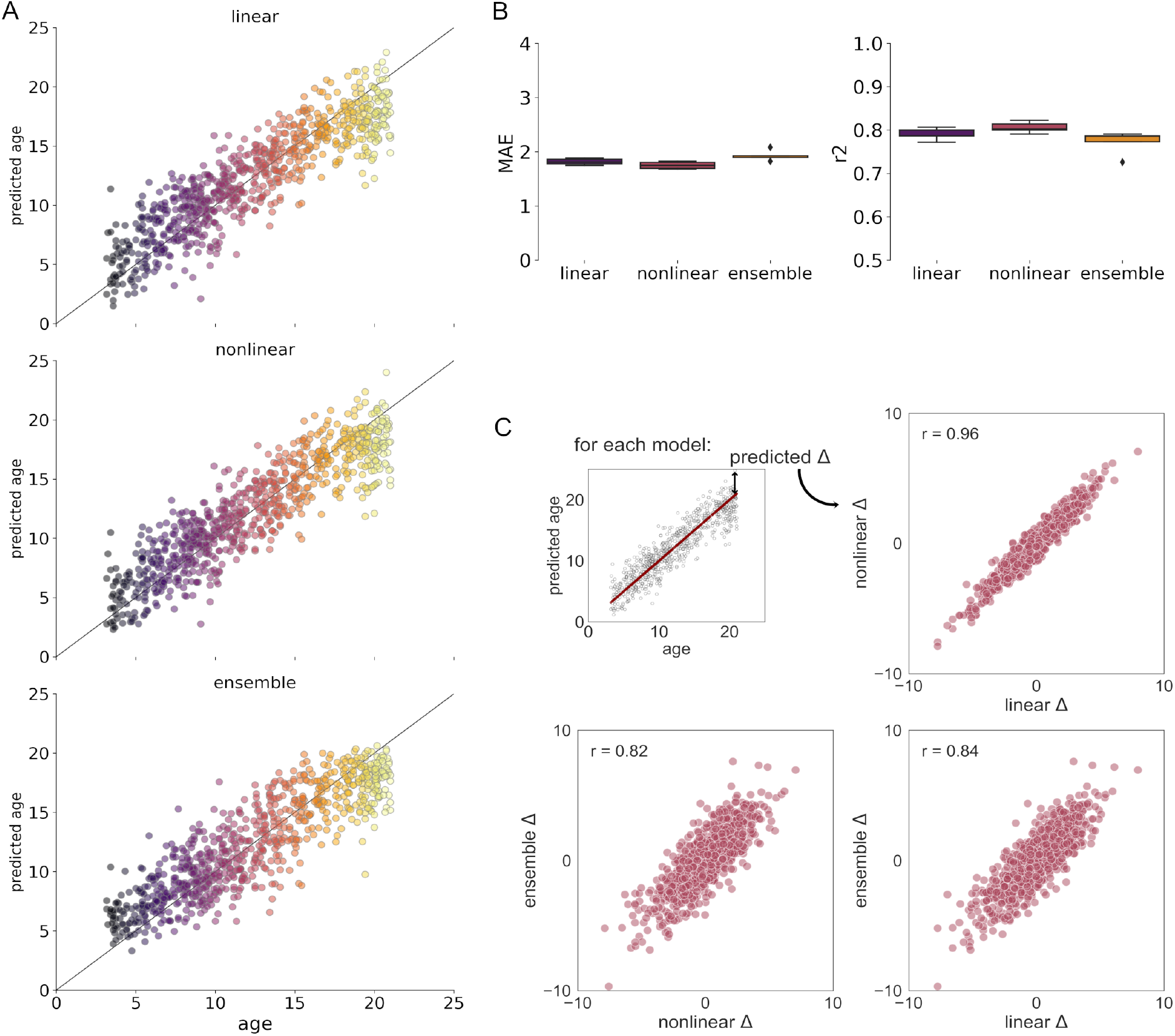
Predictions of brain age correlate across different models. A. Cross-validated predictions from each model are shown for the PING cohort. Colour indicates age and the line shows unity. B. Boxplots of median absolute error (MAE) and R^2^ score for each model, across five CV folds. C. Correlations of individual predicted age deltas (inset) between models.

Results were consistent across different scanners (Supplemental Figure S1), using alternative cortical parcellation schemes and after preprocessing with PCA (Supplemental Figure S2). Removing variance due to individual differences in mean cortical thickness and area reduced model accuracy (MAE = 2.63 [0.13], 2.63 [0.10], 2.78 [0.16]; R^2^ = 0.58 [0.06], 0.58 [0.05], 0.53 [0.07] for linear, nonlinear and ensemble models, respectively; Supplemental Figure S2), however, brain age delta estimates remained correlated with the original model estimates (linear: r = 0.81; nonlinear: r = 0.83, ensemble: r = 0.72). Across all models, brain age delta was negatively associated with age (GLM: *β*=−0.49, p<0.001; *β*=−.43, p<0.001; *β*=−0.53, p<0.001 for linear, nonlinear and ensemble models, respectively) and moderately lower in male participants (*β*=−0.15, p=0.018; *β*=−0.15, p=0.021; *β*=−0.17, p=0.006). The difference between male and female estimates of brain age error increased with age (Figure S3. Sex:age interaction: *β*=−0.14, p=0.028; *β*=−0.12, p=0.073; *β*=−0.15, p=0.017).

### Regional contributions to brain age models

We used SHAP to estimate individual-level explanations for each model prediction. For all test samples, the difference between an individual’s predicted brain age and the average prediction of an age-matched reference sample from the training set is represented as the sum of SHAP values across all cortical regions (see Methods). Thus, for a given individual, large regional SHAP values indicate where variation in cortical anatomy relative to peers contributes to a model’s prediction of brain age. Figure 3 shows the global importance of cortical features to the predictions of each model, based on mean absolute SHAP values, averaged across all subjects. For comparison, we show the model coefficients of the regularised linear model, averaged over folds (Figure 3A; Table S3). Average feature importance was correlated across models, though not exactly, with high correspondence between cortical features contributing to brain age estimation with the linear and nonlinear, GPR, models (r = 0.90). Features contributing to ensemble model predictions, while providing similarly accurate results (Figure 2), varied slightly compared to the linear and nonlinear models (r=0.60, 0.49 respectively). Results were consistent when using an alternative cortical parcellation scheme (Figure S4; linear:nonlinear r = 0.92; linear:ensemble r = 0.54; nonlinear:ensemble r = 0.44). Preprocessing with PCA increased the correlation between feature importances of the linear and nonlinear models and the ensemble model (Figure S4; linear:nonlinear r = 0.94; linear:ensemble r = 0.82; nonlinear:ensemble r = 0.71).

**Figure 3:**
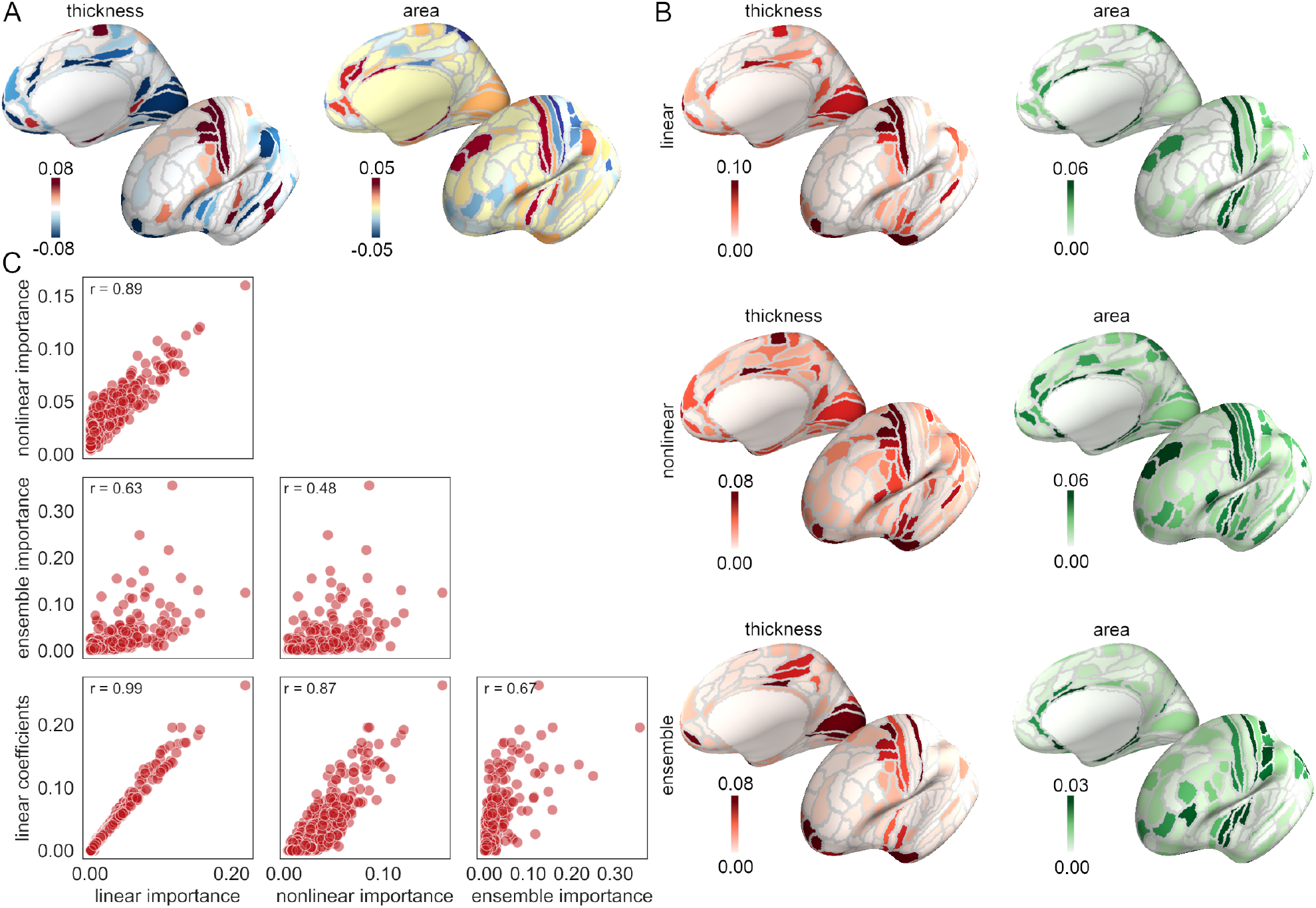
Important features correlate across models. A. Cortical surfaces display the model coefficients from Elastic Net regression averaged across five cross-validation folds (colourbar thresholded to +/− 95th centile). B. Mean absolute feature importance (SHAP values) averaged across all subjects for regional thickness (red) and area (green). Darker colours indicate greater assigned importance in the prediction of age from MRI. Colourbar shows mean importance, thresholded to the 95th centile. C. Correlations between the absolute value of the linear model coefficients and absolute feature importance averaged over subjects for each of the three brain age models. Pearson correlations are shown for each association.

Features with the largest average contribution to predictions for all models included cortical thickness of regions in the temporal pole (area TG), motor and premotor regions (frontal eye fields, primary motor cortex, area 6) and anterior frontal cortex (area 10, orbitofrontal cortex), and cortical area of regions including medial temporal lobe, insular cortex, superior temporal cortex and the temporo-parieto-occipital junction (TPOJ). The top 15 cortical thickness and area features for all models are shown in Table S4, averaged over hemispheres. The regional pattern of average feature importance remained similar across preprocessing strategies (Figure S4).

The model coefficients of the linear Elastic Net model (Table S3) reveal a similar pattern that shows thicker cortex in temporal pole (area TG), motor and supplemental motor areas (primary motor, FEF, area 6) relative to the frontal pole (areas 10, 11) and superior parietal (area 5) cortex and greater cortical area in superior frontal cortex (ventral area 6), insula (area 13, posterior insula) and motor cortex relative to somatosensory and superior parietal regions (areas 1, 7) support an older age prediction.

### Contribution of confounding variables

We found that both subject age and sex were significantly associated with brain age delta (Figure S3) and that brain age model accuracy was reduced when additionally correcting for global differences in thickness and area (Figure S2). As SHAP values are dependent on model output, and therefore brain age delta, we reasoned that anatomical variation correlated with brain age and associated with each of the confounding variables could also be represented as latent patterns present in the individual model explanations (Figure 4).

**Figure 4:**
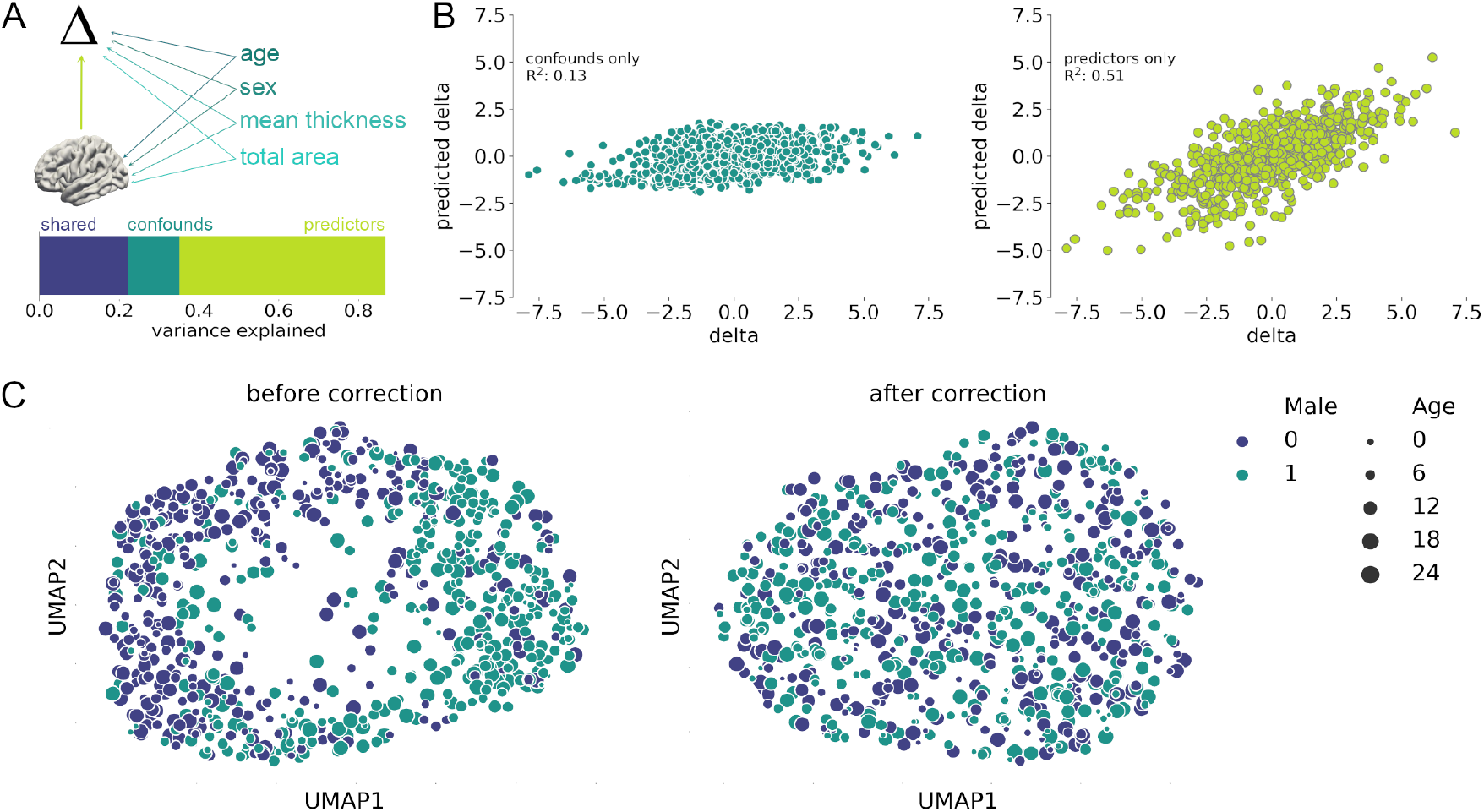
Influence of confounding variables on brain age delta. A. Top. Age, sex, mean cortical thickness and cortical surface area were identified as potential confounding variables to the association between individual brain anatomy and brain age delta. Predicted relationships between variables are shown with arrows. Associations between age, sex and mean cortical thickness and area are excluded for clarity. Bottom. Variance in brain age delta explained by shared variance across both confounding and predictor variables (dark blue), confounding variables only (teal) and predictor variables only (green). B. Semipartial correlations plots show the independent contributions of confounding (Left) and predictor (Right) variables to brain age delta. C. Two-dimensional UMAP projections of model explanations before (left) and after (right) removal of variance associated with confounding variables. Marker colour indicates sex, size indicates age.

In order to quantify the degree to which these confounding variables contribute to brain age prediction and consequently the corresponding model explanations, we performed a variance partition analysis. Following [Dinga et al., 2020], we fit a series of linear models to predict brain age delta using both confounders and model explanations. Focusing on the model with highest accuracy (GPR, see Table S5 for other models), confounding and predictor variables together accounted for 87% of variance in brain age delta (Figure 4A). Of which, variance in the model explanation data that could be also attributed to the confounding variables accounted for 22% (Figure 4A, shared). The independent contribution of confounders was 13% (Figure 4B) and, after removing variance in the explanation data associated with the confounding variables, model explanations accounted for approximately half (51%) of the variance in brain age delta (Figure 4B, right).

To illustrate the impact of confounding variables on the model explanations, we projected the *n*-subject by *p*-feature model explanations into two dimensions using a nonlinear manifold embedding technique (UMAP; [McInnes et al., 2018]). The latent variation associated with sex is apparent in Figure 4C, with male and female subjects separated along the first axis. After residualising each feature’s explanation data with respect to the set of confounding variables, this pattern is no longer evident (Figure 4C, right).

Using linear regression, we removed variance associated with the confounding variables from each model’s explanation matrix as well as brain age delta estimates before further analysis.

### Widespread regional contributions to brain age prediction

Using cross-validated, residualised model explanation data, we investigated how SHAP values depend on differences in cortical metrics during development. Figure 5A shows all regional SHAP values estimated from the nonlinear GPR model for all subjects. Regions are ordered based on the absolute value of the coefficients from the linear model, averaged over folds.

**Figure 5:**
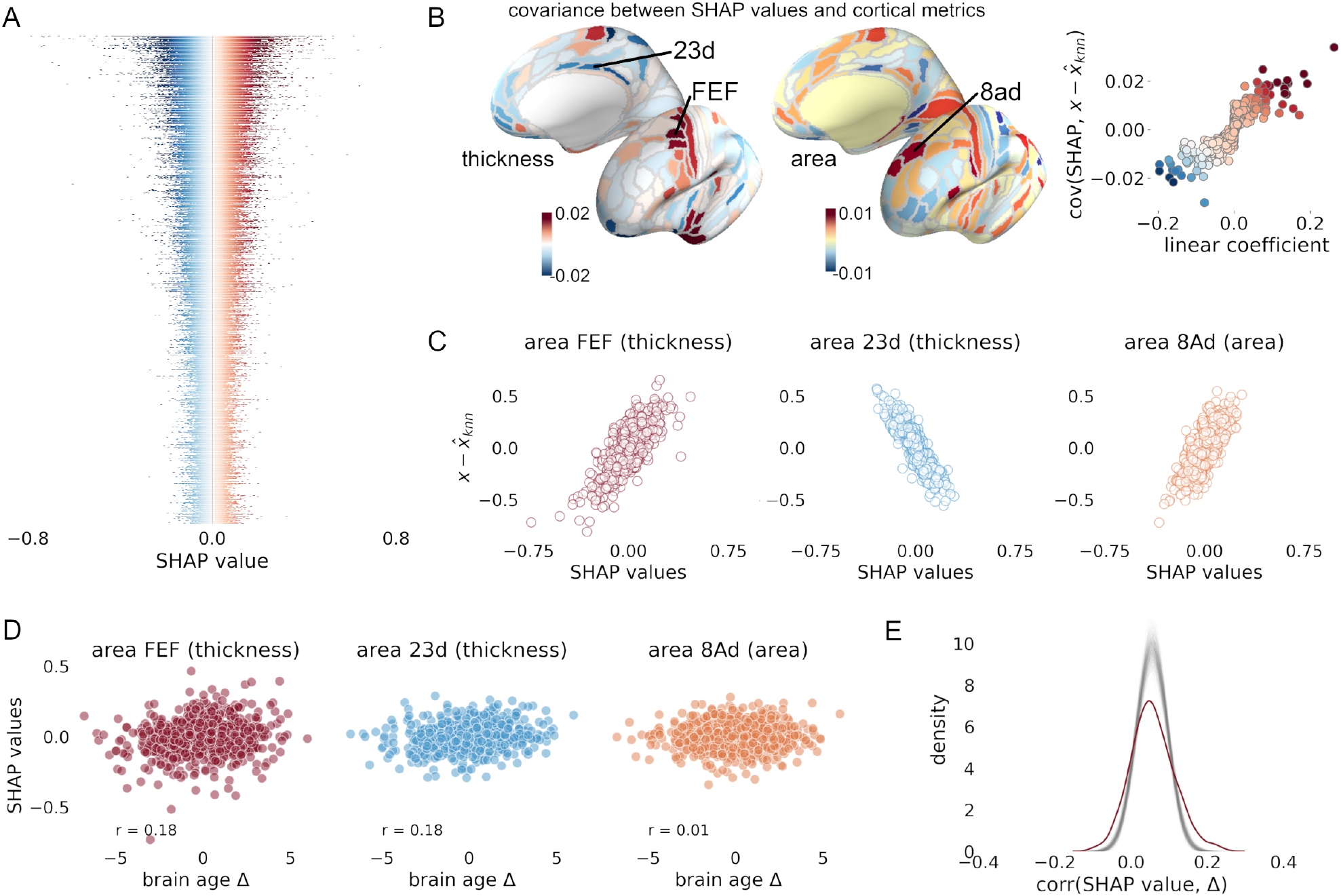
Distributed contribution of regional metrics to brain age prediction. A. SHAP summary plot of each feature, ordered by absolute magnitude of coefficients from the linear Elastic Net model, averaged over folds. Each row represents a regional metric (either thickness or area) and, in each row, each subject is represented by a marker coloured by SHAP value (negative=blue, positive=red) denoting the contribution of differences in each metric to that subject’s model prediction. B. Left, dependency of SHAP values (each row of A) on mean metric differences in each region, calculated for each feature as the difference between each subject’s value and the mean value of its *k*-nearest neighbourhood. Right, scatterplot showing the correlation between SHAP dependency and linear model coefficients. C. For three selected regions, labelled in B, the SHAP dependency on cortical metrics is shown. D. For the same three regions, the association between SHAP values and brain age delta (residualised on confounding variable) are shown. E. The distribution of correlations between SHAP values and brain age delta for all subjects, shown in red. Also shown are distribution from n=500 random surrogate maps with matched spatial autocorrelation.

We observed that, although estimated from different models, features with large contributions to individual predictions (i.e.: those with high SHAP variance across subjects) were generally those with larger model coefficients in the Elastic Net model (Figure 5A). We calculated the linear dependency of regional SHAP values on changes to cortical metrics (Figure 5B), identifying where deviation from a group of age-matched peers results in a positive or negative change in SHAP value. We found that this pattern largely recaptured the linear model coefficients and global feature importance maps shown in Figure 3 (Figure 5B, right).

However, although anatomical variation in these regions has a large relative impact on model predictions (Figure 5C), we found that the overall contribution to the final predicted brain age delta was lower (Figure 5D). Indeed, few regions exhibited larger associations between SHAP values and brain age delta than could be expected by chance (Figure 5E). This suggests that, while linear model coefficients can be used as a proxy to identify important features underlying brain age models, on an individual level the magnitude and direction of regional changes in those regions do not necessarily correlate with individual model prediction error, or brain age delta.

### Individual variation in brain age model explanations

Figure 6 shows explanations for brain age prediction using the nonlinear model in three subjects, corrected for age, sex and global cortical metrics. All three subjects were scanned at the same site and are of similar age and the same sex. Subjects *a* and *b* have similar brain age deltas, with subject *c*’s brain age predicted much lower. Regional contributions to brain age prediction varied significantly across all three subjects. Corresponding maps are shown for the linear and ensemble models in Figure S5.

**Figure 6:**
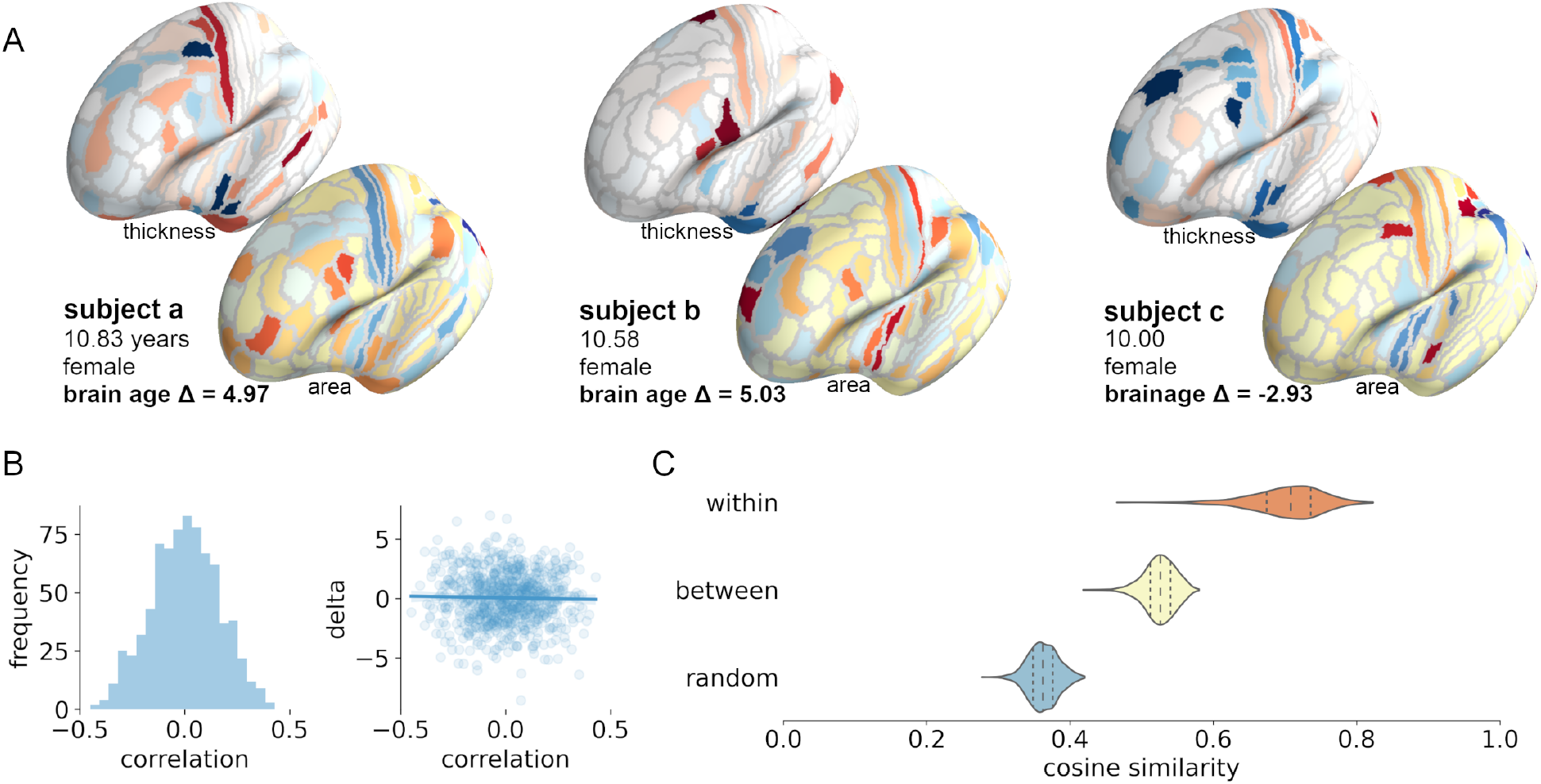
Individual variation in features explaining brain age prediction error. . A. Individual model explanations for the GPR model are shown to the same scale (−.15 to .15) for three subjects of similar age on the inflated cortical surface, after removing variance due to age, sex and global cortical metrics. B. Left, distribution of correlations between individual model explanations and the linear model coefficients. Right, association between correlations and residualised brain age delta. C. Distribution of cosine similarities between absolute value model explanations estimated within the same subject (across training/test folds), between different subjects, and between explanations and random surrogate maps (n=500). The absolute value was used to account for potential anti-correlated feature values in subjects with opposing brain age deltas.

Overall, model explanations for a given subject were similar when using different models. For each subject, we quantified the similarity of model explanations across all three models, we found mean (S.D) within-subject cosine similarity of 0.79 (0.048) between individual’s linear and nonlinear model explanations, 0.47 (0.082) between linear and ensemble explanations, and 0.38 (0.071) between nonlinear and ensemble explanations. This compared to mean between-subject similarities across models of 0.00 (0.094), 0.00 (0.102) and 0.00 (0.073), respectively (Figure S6).

While consistent across models, we found that, on an individual level, model explanations were not generally correlated with the coefficients derived from the regularised linear model. Using explanations from the nonlinear GPR model, we found that correlations ranged between approximately −0.5 and 0.5 and were not associated with brain age delta (Figure 6B). This observation was largely consistent across other models (Figure S7). SHAP assigns importance using a sparse local regression where the number of contributing features is set via ℓ_1_ regularisation. We repeated our analysis specifying fewer features (maximum *n*=50) to model each subject’s brain age predictions (Figure S8). Overall, the results were similar, however using the reduced set of features we noted a moderate positive correlation between brain age delta and the similarity between subject-specific nonlinear model explanations and the linear model coefficients (*r* = 0.31; Figure S8F).

Finally, we tested the degree of spatial agreement between different individual’s explanations for a given model. For each model, we took advantage of the multiple versions trained during cross-validation to estimate within-subject similarity of model explanations across training folds. Using absolute SHAP values to account for potential anti-correlation of regional explanations in subjects with opposing brain age deltas, we then calculated mean similarity between each subject’s maps and that of every other in the cohort. Overall, we found high similarity between model explanations in the same subject estimated across training folds (linear model: mean [S.D.] = 0.77 [0.043]; nonlinear: 0.70 [0.048]; ensemble: 0.61 [0.104]), compared to the similarity between different subject’s regional explanations for the same model (linear model: 0.54 [0.023]; nonlinear: 0.52 [0.022]; 0.49 [0.058]. To test whether the degree of spatial similarity was greater than expected by chance (i.e.: if no common structure was shared across individual model explanations), we constructed a null distribution, measuring the similarity between each subject’s model explanations and a set of randomly generated maps with matched intensity distributions and spatial autocorrelation. We found significantly lower similarities compared to within- and between-subject comparisons (linear: 0.19 [0.013]; nonlinear: 0.37 [0.021]; ensemble: 0.15 [0.019]) (Figure 6C). In contrast, repeating this analysis specifying fewer explanatory features in SHAP, we found that while within-subject similarity (nonlinear: mean [S.D.] = 0.40 [0.102]) remained greater than between subjects (mean [S.D.] = 0.31 [0.047]), within-subject model explanations were highly variable across folds (range: 0.06 - 0.68; Fig S9G).

### No associations between brain age delta and cognition

Finally, in a subset of the cohort with available data (n=573), we tested the association between cross-validated estimates of brain age delta and composite measures of cognitive performance from the NIH Cognitive Toolbox. Age and sex accounted for between 41% and 66% of variance in each of the NTCB scores (Table S6). After residualising each with respect to these factors, we projected the scores onto the two largest principal components, that explained 33% and 21% of the remaining variance, respectively. The loading of each scale onto the principal components (PCs) are shown in Table S7. Using linear regression, we found no significant associations between brain age delta estimates from any of the three models and cognitive performance (Table S8).

## Discussion

In this study, we examined brain age modelling approaches in a cohort of typically developing children. We find that brain age estimates are relatively robust and highly correlated across different models and parameter settings. We show that age and sex are significantly associated with brain age predictions, together explaining around 35% of variance in brain age prediction in this cohort. Accounting for this, we find that regional cortical changes apparent in development are important factors in brain age prediction, on average. However, we demonstrate that, on an individual basis, the contribution of changes in these regions to a final model prediction varies significantly across subjects, even when age, sex and brain age delta are all similar. Finally, we didn’t find any evidence that brain age error is associated with cognitive performance in this typically-developing sample.

Brain age modelling has recently become a popular approach to modelling variation in brain development and aging in both healthy and clinical populations [Cole et al., 2018; Cole and Franke, 2017; Franke and Gaser, 2012; Franke and Gaser, 2019; Jiang et al., 2020; Kaufmann et al., 2019; Liem et al., 2017; Niu et al., 2020]. Previously, studies have shown that although model performance can vary, different model architectures can achieve similarly accurate age predictions [Cole et al., 2017; Franke et al., 2010; Jiang et al., 2020; Schulz et al., 2020b]. Here, we observed similar performance across three different types of models: regularised linear regression, Gaussian process regression and tree-based ensemble. All models achieved median absolute errors of less than 2 years, in line with previous examples in this and other paediatric cohorts [Ball et al., 2017; Brown et al., 2012; Khundrakpam et al., 2015; Lewis et al., 2018].

Further, we found that individual cross-validated estimates of brain age were highly correlated across model types (between *r* = 0.82 and 0.94). Consistent with previous reports, we also find that brain age estimates were not strongly affected by processing parameters including cortical parcellation scheme [Khundrakpam et al., 2015], and dimension reduction with PCA [Franke et al., 2010].

As others have reported, brain age delta varied across the cohort as a function of age [Butler et al., 2020; Le et al., 2018a; Liang et al., 2019; Smith et al., 2019]. This is commonly attributed to a ‘regression towards the mean’ effect and can be corrected through linear methods *post hoc* [Beheshti et al., 2019; Le et al., 2018a; Smith et al., 2019], or by including age as an additional covariate in subsequent analyses [Le et al., 2018a], though recent reports have urged caution in the use and interpretation of brain age deltas (corrected or otherwise) in subsequent models [Butler et al., 2020]. To avoid potentially inflating model performance, we have reported accuracies based on the raw prediction error, without additional correction for age bias. We also found that participant sex was associated with significant differences in brain age delta. Despite small mean differences and considerable overlap in brain neuroanatomy [Ritchie et al., 2018], multivariate statistical methods are able to robustly classify male and female brains with high accuracy [Anderson et al., 2019; Ball et al., 2017; Chekroud et al., 2016; Peng et al., 2020]. Other studies have reported this effect in brain age models [Brouwer et al., 2020; Cole et al., 2018], or have fit separate models to male and female samples to account for it [Erus et al., 2015; Goyal et al., 2019; Han et al., 2020]. Here, we find that brain age delta was on average higher in females than males (i.e.: brain-predicted age was older), and that this difference increased with age. This may reflect accelerated brain maturation in females in mid-to-late adolescence, coinciding with timing of pubertal development [Brouwer et al., 2020; Gennatas et al., 2017].

Together with measures of global cortical metrics (mean cortical thickness and total surface area), we found that age and sex accounted for approximately half of the variance in brain age delta. Interestingly, neither mean cortical thickness or total surface area correlated with brain age delta (*r*~0.01-0.06 across all three models), suggesting that the observed decrease in performance after correcting regional brain features for global metrics reflects removal of variance associated with mean differences between sexes. We chose to model both sexes together, rather than train separate models, in order to enable clearer interpretation of explanations derived from the same model. An interesting future direction may be to explore if feature-level explanations of separate brain age models vary systematically across male and female samples.

PING is a multi-site study combining information from several MRI scanners. Brain age prediction is generally poorer when no samples from the same site or scanner are included in the model training data set [Cole et al., 2017; Franke et al., 2010; Liem et al., 2017]. Using a multi-model ensemble model in an adult cohort aged 19 to 82 years, Liem et al. reported an 86% increase in MAE from 4.29 to 8.02 years when predicting samples from a new site unseen by the model. Introducing some training samples from the new site reduced MAE to 6.93 years [Liem et al., 2017]. In the PING cohort, Khundrakpam et al. reported much smaller increases in cross-validated MAE of around 0.03 - 0.08 years, when applying brain age models to unseen sites [Khundrakpam et al., 2015]. This likely reflects the smaller age range and the effort made to ensure harmonised acquisition protocols across sites and scanners in the PING cohort [Jernigan et al., 2016]. Here, we applied ComBat to account for linear shifts in feature distributions across sites [Fortin et al., 2018] and performed a stratified 5-fold cross-validation, balancing each fold by site, to ensure stable brain age predictions across sites (Fig S1). As methods such as SHAP explain the model, rather than the data, lower model accuracy from a site-specific bias in out-of-sample data would be reflected in the model explanations, with site-specific differences represented as a latent pattern present in the model explanation data matrix. This would make comparisons in model explanations across scanners in a multisite study challenging. By applying a site-correction beforehand, and ensuring stratified samples in cross-validation, we were able to focus on patterns of individual variation rather than site bias.

Using SHAP, we found that anatomical variance in a common set of regions drives model predictions of age, regardless of model type. On average, these regions were similarly weighted in a regularised linear model. However, we found that individual level explanations, while relatively stable across model types and training folds, do not necessarily correlate with overall feature importance. As such, the strength of contribution of each region to the final model prediction varied significantly across subjects, with limited overlap between subjects, even when matched for age, sex and brain age delta. Regularised linear models can achieve accuracies on par with more complex, nonlinear or deep examples [He et al., 2020; Schulz et al., 2020b]. A main benefit of linear approaches is the interpretability offered by the model coefficients, each representing the weighted contribution of a feature to the final model prediction. Linear model coefficients are often used to identify important features in brain age modelling [Aycheh et al., 2018; Khundrakpam et al., 2015; Lewis et al., 2018; Zhai and Li, 2019]. Using Elastic Net regression, we find that age prediction is based on the relative thickness of primary and secondary motor regions relative to primary visual cortex, and lateral parietal cortex, as well as the area of primary motor and superior frontal cortex relative to primary sensory cortex. This pattern reflects developmental growth patterns of the cortex in childhood and adolescence, with thinner cortex and greater area in fronto-parietal association cortex relative to primary somato-motor regions in older subjects [Ball et al., 2020; Mills et al., 2016; Raznahan et al., 2011; Reardon et al., 2018; Shaw et al., 2006; Tamnes et al., 2017].

We found that regions with a high weight in the linear model were, on average, assigned high importance by SHAP for individual model decisions. Overall, we found subject explanations of brain age to be stable across model types and model instances trained using different subsamples during cross-validation. However, we found that individual explanations did not necessarily correlate with the linear model coefficients. For a linear model, this is expected given the relationship between brain age delta, SHAP values, model coefficients and an individual’s data: 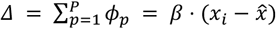, where regional SHAP values relate to both the model coefficients, *β*, and regional neuroanatomical differences between a given subject relative to a reference baseline, generally the sample mean 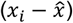. In this case, the model coefficients accurately represent a model of age as a function of brain anatomy across the whole group but, for many brain age studies, we are interested not in the regional patterns that underlie an accurate age prediction, but in the individual differences that underlie error in that prediction. In this case, we found that even when assigned a high SHAP value, the overall contribution of a given region to final brain age predictions were generally small. This indicates that brain age delta is best explained by widespread changes across multiple regions, the pattern of which can vary significantly across subjects. We note that, by construction, SHAP values are spread across multiple explanatory features when fitting the local linear regression to model outputs. By applying ℓ_1_ regularisation to the original model features, a subset is selected in order to aid interpretation. We initially set this regularisation to select, at most, 50% of cortical features. We repeated our analysis using a much higher threshold, selecting at most 7% (*n*=50), and observed largely similar results. In contrast to the main results, however, we found that within-subject model explanations were much less stable when built using fewer features. We believe that the limited association between model coefficients and individual feature importance is an important consideration for future brain age studies.

As with previous studies in this cohort, we did not find strong evidence of association between brain age delta and cognitive performance, after accounting for age and sex [Ball et al., 2017; Khundrakpam et al., 2015]. Other studies have reported small to moderate associations between brain age and measures of cognitive performance during typical development [Erus et al., 2015; Lewis et al., 2018], or associations between brain age estimates and symptoms of psychopathology or neurodevelopmental disorders [Cropley et al., 2020; Tunç et al., 2019]. Our findings do not preclude the detection of brain age-behaviour associations in typically-developing samples, others have employed multi-modal approaches to successfully predict brain age from brain structure and function, additionally using measures of tissue microstructure from diffusion MRI and subcortical metrics rather than just cortical thickness and area [Erus et al., 2015; Lewis et al., 2019; Liem et al., 2017]. In typical samples, however, associations are generally small and likely require much larger sample sizes to reliably estimate [Helmer et al., 2020; Kharabian Masouleh et al., 2019; Marek et al., 2020]. We agree with recent suggestions that model explanations may prove useful in dissecting or stratifying brain age-behavior associations, particularly in clinical populations where individual variation in neuroanatomy may reflect differential patterns of neurodevelopment in heterogeneous populations [Schulz et al., 2020a].

In this study, we employ kernelSHAP to generate individual model explanations. SHAP generalises several existing model explanation methods and can be applied across different model types [Lundberg and Lee, 2017]. Exact methods exist for linear models (linearSHAP, for models without regularisation) and tree-based models (treeSHAP), such as XGBoost [Lundberg et al., 2020]. One key feature is the ability to produce local explanations on a subject-by-subject basis compared with other approaches that provide global estimates of feature importance over the full sample, e.g.: permutation feature importance [Breiman, 2001]. A key assumption of the kernelSHAP method is feature independence, this can result in inaccurate attribution of importance when features are highly correlated [Aas et al., 2020]. As feature dependence can be accommodated by treeSHAP, we tested the similarity between kernelSHAP explanations and treeSHAP explanations with and without feature independence for the ensemble model predictions (Figure S9). We found high correspondence across individuals between the approximated (kernelSHAP) and exact (treeSHAP) model explanations. Recent releases of kernel SHAP methods that can account for feature dependence should prove useful for model-agnostic explanation of machine learning models in neuroimaging [Aas et al., 2020].

Another key consideration is that SHAP explains the model, not the data [Lundberg and Lee, 2017]. Feature importance can be viewed as the distribution of prediction error across model features. Features that aren’t used by a model will have lower importance, regardless of any underlying associations with the variable of interest. Although feature importance was generally correlated across model types, this is evident in the ensemble model, where global feature importance was less similar than between linear and GPR models. A poorly fitting model will also have high prediction error and SHAP values may not necessarily reflect the true importance of a given feature to the modelled process. Here, we achieved good model performance across all model types and, by using an age-matched reference sample, we focused on explanations of age prediction error relative to age-matched peers. While increasing model accuracy would generally be considered the main aim of any modelling approach, by its derivation, brain age delta is model error and reducing it too much can obfuscate apparent neuroanatomical differences between clinical groups [Bashyam et al., 2020].

While we show that model explanations are consistent within-subject, we are not able to determine if this remains so over time due to the cross-sectional study. Test-retest and longitudinal studies have shown that individual estimates of brain age delta are stable over both short (weeks) and long-term (years) periods [Brouwer et al., 2020; Cole et al., 2017]. Future studies may employ model explanation methods to determine if feature importance to individual predictions also remains stable.

We find that brain age estimates in typical development are generally robust to model and preprocessing choice. Brain age delta is associated with both age and sex and reflects developmental remodelling in the cortex. However, the importance of a given region to prediction of brain age varies significantly across subjects even when matched for age, sex and brain age error. While linear models generally offer good performance for datasets commonly seen in larger neuroimaging studies and provide clearly interpretable estimates of average model importance in the feature weights, we recommend the use of explanation methods to aid interpretation of complex or nonlinear models, particularly when individual variation is a key property of interest. These methods are generally stable of model choices, though require additional parameter choices that depend on the type of model used. We reiterate that interpretation of model explanations depends both on the data and the model fit.

## Data availability

Data from the PING study are made available via the NIMH Data Archive (https://nda.nih.gov/edit_collection.html?id=2607).

Python code supporting this study is available at: https://github.com/garedaba/brainAges

## Acknowledgements

This research was conducted within the Developmental Imaging research group, Murdoch Childrens Research Institute and the Children’s MRI Centre, Royal Children’s Hospital, Melbourne, Victoria. It was supported by the Murdoch Childrens Research Institute, the Royal Children’s Hospital, Department of Paediatrics, The University of Melbourne and the Victorian Government’s Operational Infrastructure Support Program. The project was generously supported by RCH1000, a unique arm of The Royal Children’s Hospital Foundation devoted to raising funds for research at The Royal Children’s Hospital.

Data and/or research tools used in the preparation of this manuscript can be obtained from the controlled access datasets distributed from the NIMH-supported Research Domain Criteria Database (RDoCdb). RDoCdb is a collaborative informatics system created by the National Institute of Mental Health to store and share data resulting from grants funded through the Research Domain Criteria (RDoC) project.

## Notes

### Competing Interest Statement

The authors have declared no competing interest.

